# Fibrinolysis influences SARS-CoV-2 infection in ciliated cells

**DOI:** 10.1101/2021.01.07.425801

**Authors:** Yapeng Hou, Yan Ding, Hongguang Nie, Hong-Long Ji

## Abstract

Rapid spread of COVID-19 has caused an unprecedented pandemic worldwide, and an inserted furin site in SARS-CoV-2 spike protein (S) may account for increased transmissibility. Plasmin, and other host proteases, may cleave the furin site of SARS-CoV-2 S protein and γ subunits of epithelial sodium channels (γ ENaC), resulting in an increment in virus infectivity and channel activity. As for the importance of ENaC in the regulation of airway surface and alveolar fluid homeostasis, whether SARS-CoV-2 will share and strengthen the cleavage network with ENaC proteins at the single-cell level is urgently worthy of consideration. To address this issue, we analyzed single-cell RNA sequence (scRNA-seq) datasets, and found the PLAU (encoding urokinase plasminogen activator), SCNN1G (γENaC), and ACE2 (SARS-CoV-2 receptor) were co-expressed in alveolar epithelial, basal, club, and ciliated epithelial cells. The relative expression level of PLAU, TMPRSS2, and ACE2 were significantly upregulated in severe COVID-19 patients and SARS-CoV-2 infected cell lines using Seurat and DESeq2 R packages. Moreover, the increments in PLAU, FURIN, TMPRSS2, and ACE2 were predominately observed in different epithelial cells and leukocytes. Accordingly, SARS-CoV-2 may share and strengthen the ENaC fibrinolytic proteases network in ACE2 positive airway and alveolar epithelial cells, which may expedite virus infusion into the susceptible cells and bring about ENaC associated edematous respiratory condition.

## Introduction

The SARS-CoV-2 infection leads to COVID-19 with pathogenesis and clinical features similar to those of SARS and shares the same receptor, angiotensin-converting enzyme 2 (ACE2), with SARS-CoV to enter host cells (Zhou et al. 2020, Li and Zheng 2020). By comparison, the transmission ability of SARS-CoV-2 is much stronger than that of SARS-CoV, owning to diverse affinity to ACE2 (Wrapp and Wang 2020). The fusion capacity of coronavirus *via* the spike protein (S protein) determines infectivity (Wrapp and Wang 2020, Kam et al. 2009b). Highly virulent avian and human influenza viruses bearing a furin site (RxxR) in the haemagglutinin have been described (Coutard et al. 2020). Cleavage of the furin site enhances the entry ability of Ebola, HIV, and influenza viruses into host cells (Claas et al. 1998). Consisting of receptor-binding (S1) and fusion domains (S2), coronavirus S protein needs to be primed through the cleavage at S1/S2 site and S2’ site for membrane fusion (Jaimes et al. 2020, Huggins 2020). The newly inserted furin site in SARS-CoV-2 S protein significantly facilitated the membrane fusion, leading to enhanced virulence and infectivity (Xia et al. 2020, Wang, Qiu, et al. 2020).

Plasmin cleaves the furin site in SARS-CoV S protein (Kam et al. 2009b), which is upregulated in the vulnerable populations of COVID-19 (Ji et al. 2020). However, whether plasmin cleaves the newly inserted furin site in the SARS-CoV-2 S protein remains obscure. Plasmin cleaves the furin site of human γ subunit of epithelial sodium channels (γENaC) as demonstrated by LC-MS and functional assays (Zhao, Ali, and Nie 2020, Sheng et al. 2006). Very recently, it has been proposed that the global pandemic of COVID-19 may partially be driven by the targeted mimicry of ENaC α subunit by SARS-CoV-2 (Gentzsch and Rossier 2020, Muhanna et al. 2020). ENaC are located at the apical side of the airway and alveolar cells, acting as a critical system to maintain the homeostasis of airway surface and alveolar fluid homeostasis (Ji et al. 2006, Matalon, Bartoszewski, and Collawn 2015). The luminal fluid is required for keeping normal ciliary beating to expel inhaled pathogens, allergens, and pollutants and for migration of immune cells that release pro-inflammatory cytokines and chemokines (Hou et al. 2019a). The plasmin family and ACE2 are expressed in the respiratory epithelium (Nie et al. 2009, Hanukoglu and Hanukoglu 2016, Kam et al. 2009a). However, if the plasmin system and ENaC are involved in the fusion of SARS-CoV-2 into host cells is unknown.

This study aims to determine whether PLAU, SCNN1G, and ACE2 are co-expressed in the airway and lung epithelial cells and whether SARS-CoV-2 infection alters their expression at the single-cell level. We found that these genes, especially the PLAU was significantly upregulated in epithelial cells of severe/moderate COVID-19 patients and SARS-CoV-2 infected cell lines, mainly owning to ciliated cells. We conclude that the most susceptible cells for SARS-CoV-2 infection could be the ones co-expressing these genes and sharing plasmin-mediated cleavage.

## Results

### Furin sites are identified in both virus and host γENaC proteins

A furin site was located at the S proteins of SARS-CoV-2 from Arginine-683 to Serine-687 (RRAR|S), and similar site was also seen in the S protein of HCoV-OC43, MERS, and HCoV-HKU1 coronavirus (**Fig. 1A**). In addition, the highly conserved RxxR motif existed in the hemagglutinin protein of influenza H3N2, Herpes, Ebola, HIV, Dengue, hepatitis B, West Nile, Marburg, Zika, Epstein-Barr, and respiratory syncytial virus (RSV). The furin site (RKRR|E) was found in the gating relief of inhibition by proteolysis (GRIP) domain of the extracellular loop of the mouse, rat, and human γENaC (**Fig. 1B**). The similarity of these furin sites is 40-80%.

**Figure 1.**
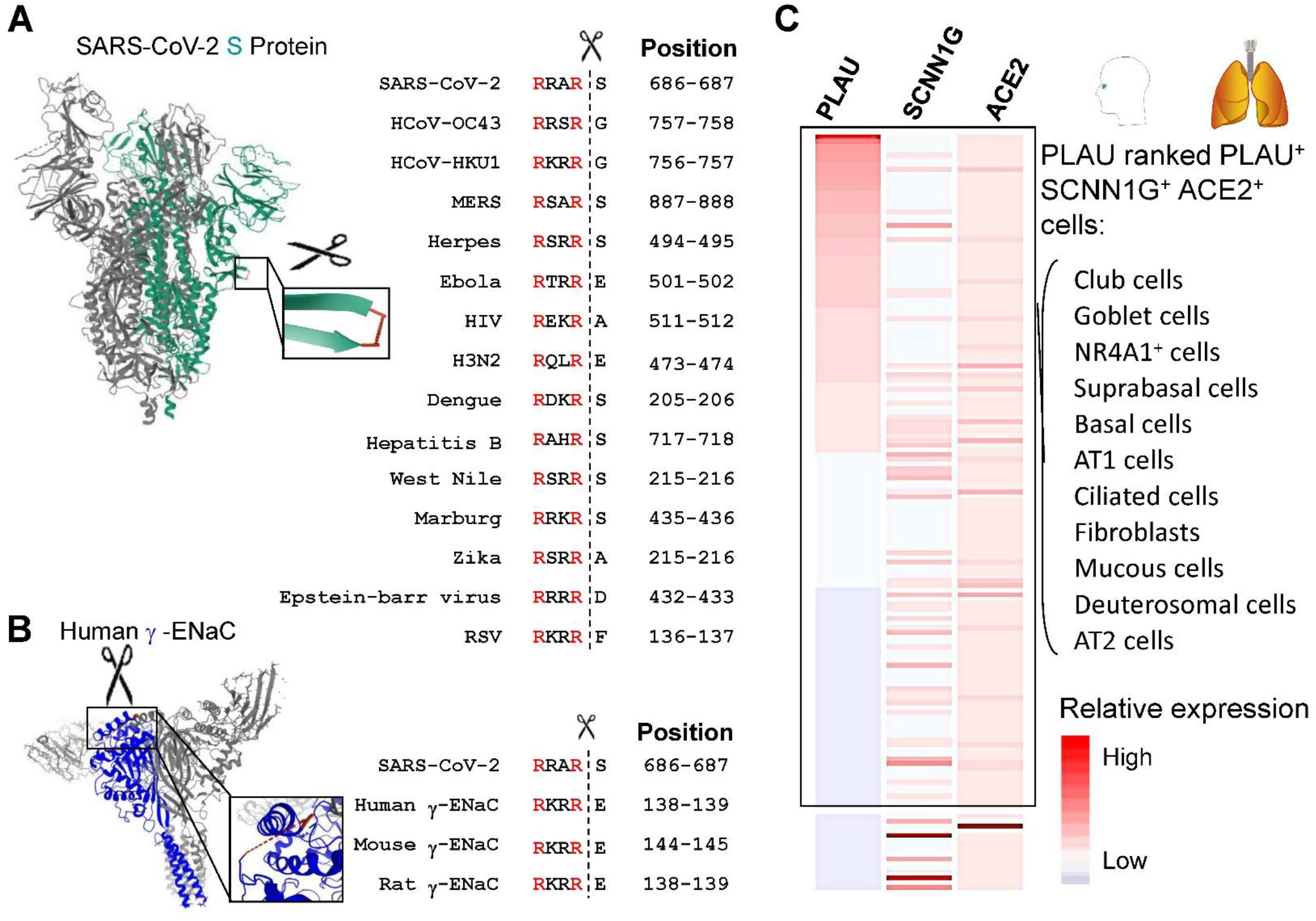
Targeted molecular mimicry by SARS-CoV-2 of human γENaC and profiling ACE2-SCNN1G-PLAU/PLAT co-expression. **(A)** The cartoon showed the S-protein of SARS-CoV-2 (PDB ID: 6X2A), which was highlighted in green. The S1/S2 cleavage site required for the activation of SARS-CoV-2 was enlarged and highlighted in red. Furin/plasmin cleavage sites of common human viruses were shown in a box. **(B)** The cartoon represents the human γENaC protein (PDB ID: 6BQN), which was highlighted in green. Furin/plasmin cleavage site was enlarged and highlighted in red. The cleavage sites of γENaC in other species were shown in a box. **(C)** The single-cell transcriptomic co-expression of ACE2, SCNN1G (γENaC), and PLAU was summarized. The heatmap depicted the mean relative expression of each gene across the identified cell populations. The cell types were ranked based on decreasing expression of PLAU. The box highlighted the ACE2, SCNN1G (γENaC), and PLAU co-expressing cell types in the human respiratory system.

### Respiratory cells co-express PLAU, SCNN1G, and ACE2

To identify subpopulations of cells co-expressing PLAU, SCNN1G, and ACE2, we analyzed 11 scRNA-seq datasets by nferX scRNA-seq platform (https://academia.nferx.com/) (**Supplementary Table 1**). All three genes were co-expressed in the following cells ranked by the expression level of PLAU from high to low: club cells, goblets, basal cells, AT1 cells, ciliated cells, fibroblasts, mucous cells, deuterosomal cells, and AT2 cells (**Fig. 1C**), which were supported by previous studies (Sungnak et al. 2020, Wang et al. 2008, Hanukoglu and Hanukoglu 2016). These results suggest that these cell populations co-expressing PLAU-γENaC-ACE2 may be more susceptible to the SARS-CoV-2 infection compared with others. In addition, the top ten ranked cell sub-populations expressing PLAU, SCNN1G, or ACE2 alone were listed in **Supplementary Table 2**. To compare the transcript of the proteases in different lung epithelial cells, we analyzed the lung dataset from Gene Expression Omnibus (GEO) by Seurat, and the cells were annotated by their specific markers (**Supplementary Fig. 1A**). The data showed that all these proteases were expressed in AT2 cells, including PLAU, FURIN, PRSS3 (Trypsin), ELANE (Elastase), PRTN3 (Myeloblastin), CELA1 (Elastase-1), CELA2A (Elastase-2A), CTRC (Chymotrypsin-C), TMPRSS4 (Transmembrane protease serine 4), and TMPRSS2 (Transmembrane protease serine 2) (**Supplementary Fig. 1B**). In AT2 cells, the proteases expression level in order is: TMPRSS2 > FURIN > TMPRSS4 > PLAU > CELA1 > ELANE > PRSS3 > PRTN3 > CTRC > CLEA2A. For PLAU, the high to low order is Basal > Club > Ciliated > AT1 > AT2.

The expression levels of proteases (PLAU, FURIN, TMPRSS2, PLG), ACE2, and SCNN1G in 11 cell types co-expressing ACE2, SCNN1G, and PLAU were compared in **Fig. 2**. The club cells showed the highest expression level of PLAU, and the ACE2, SCNN1G, TMPRSS2, FURIN, and PLG showed a higher expression level in club cells compared with other cell types. Of note, the ciliated cell was the second and seventh highest expression cell type of PLAU and ACE2, respectively.

**Figure 2.**
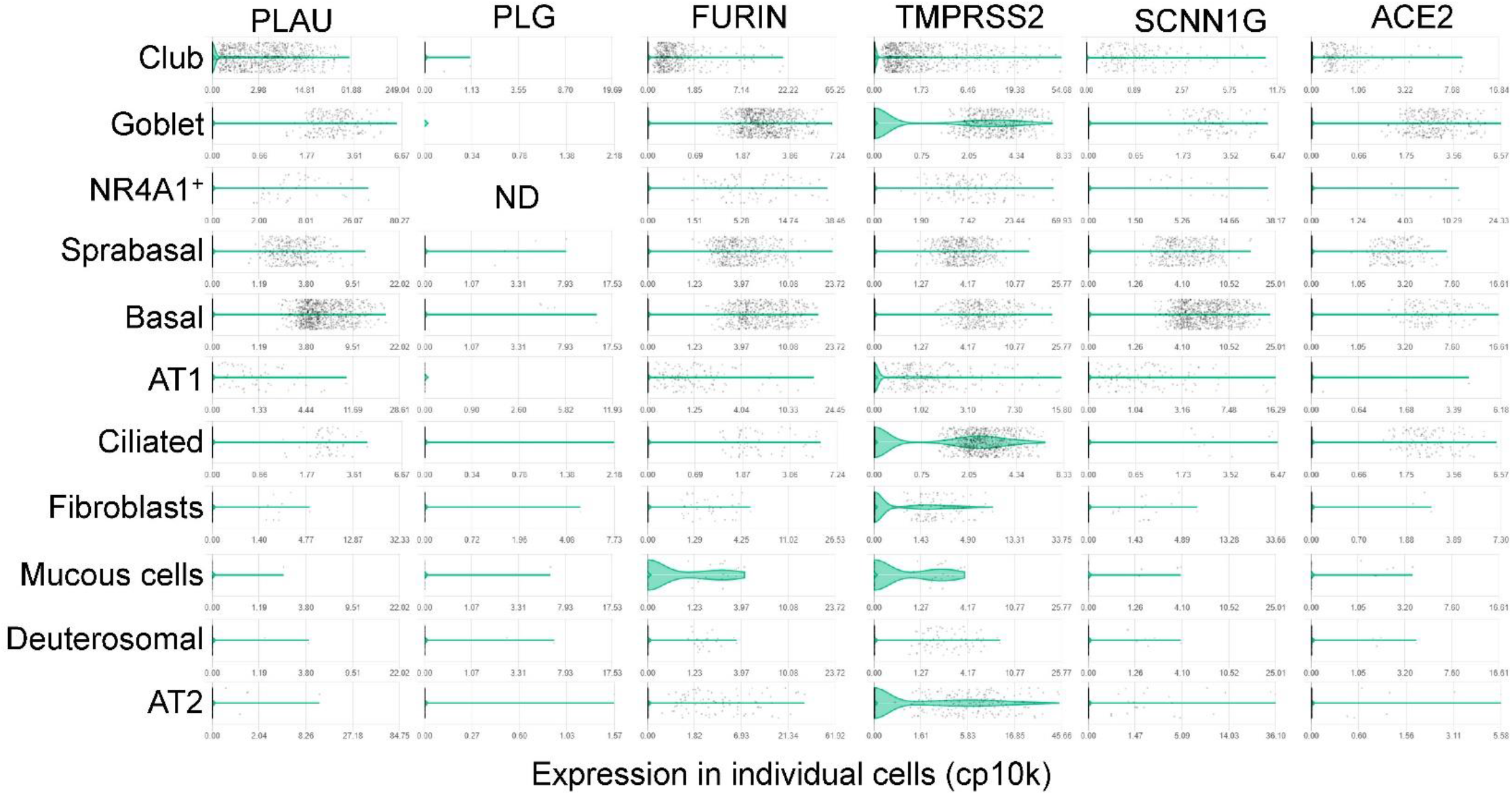
Expression of proteases, γENaC, and ACE2 in the human respiratory system. Violin plots showing the expression level of PLAU, PLG, FURIN, TMPRSS2, and SCNN1G in nferX platform.

### Expression levels of PLAU, SCNN1G, and ACE2 in SARS-CoV-2 infection

To detect the potential changes in the cell populations that co-express PLAU, SCNN1G, and ACE2, we analyzed the scRNA-seq datasets of bronchoalveolar lavage fluid (BALF) cells, which are mainly composed of epithelial cells and leukocytes. There were three groups to be studied: 4 healthy controls, 3 moderate, and 6 severe COVID-19 patients. The expression level and the percentage of total cells expressing PLAU and FURIN were significantly upregulated in the severe group compared with controls (*P* < 0.001), as well as the expression levels of ACE2, TMPRSS2, SCNN1G, and PLG were also slightly upregulated (**Fig. 3A and B**).

The expression levels of PLAU, Furin, TMPRSS2, and ACE2 and the number of cells were profiled in **Fig. 4A**. The data showed that these genes were upregulated in COVID-19 patients, and the number of cells expressing these upregulated genes almost increased in a severity-dependent manner. PLAU was significantly elevated in severe group (*P* < 0.001), and the other genes also showed an increasing trend (**Fig. 4B**).

**Figure 3.**
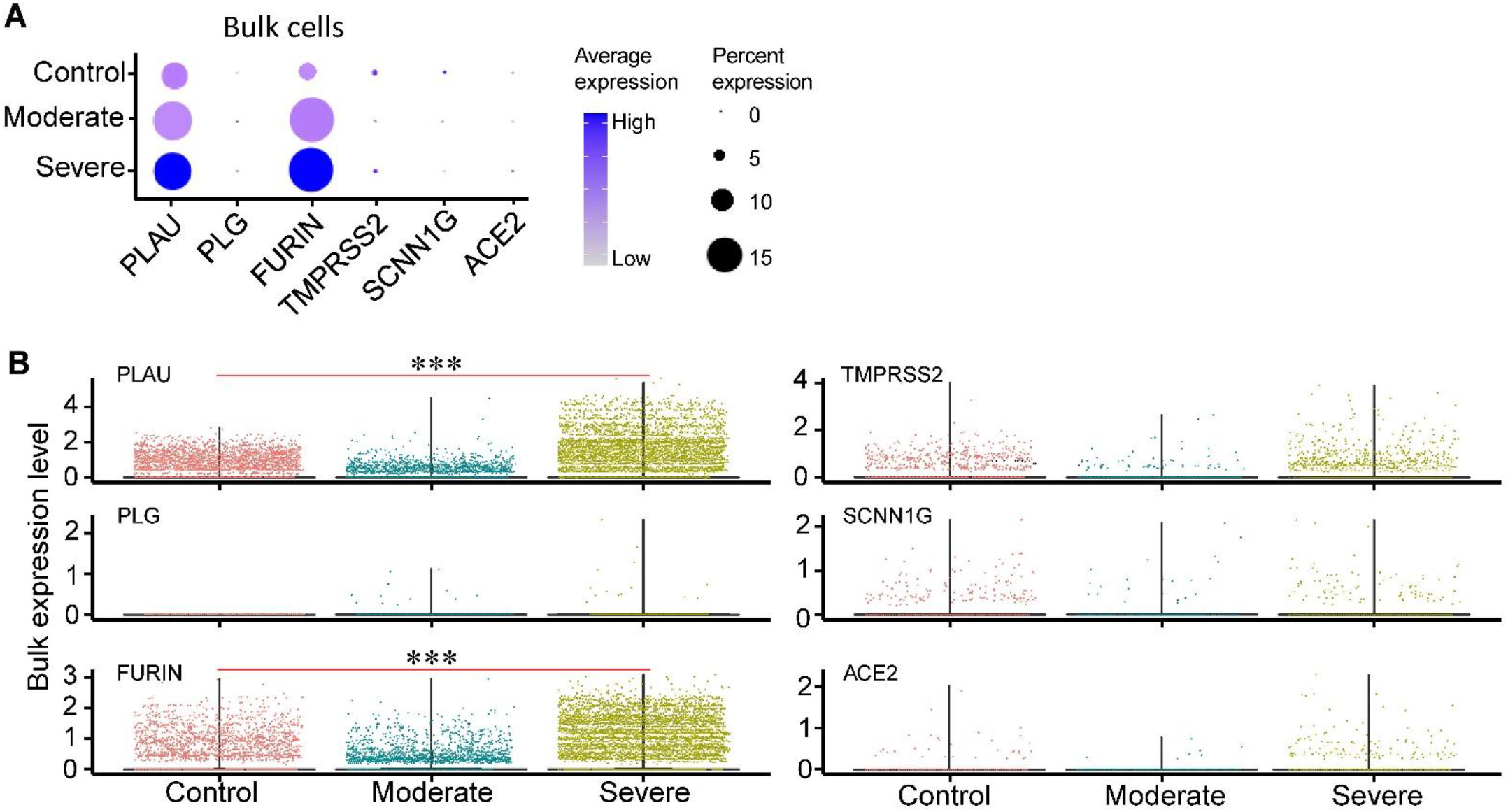
Overall expression levels of proteases, ACE2, and SCNN1G in BALF bulk cells of COVID-19 patients. **(A)** Bubble plot of proteases, ACE2, and SCNN1G in BALFs of COVID-19 patients. The size of the dots indicateed the proportion of cells in the respective cell type having a greater-than-zero expression of these genes, while the color indicated the mean expression of these genes. **(B)** The gene expression levels of proteases, ACE2, and SCNN1G from health controls (n = 4), moderate cases (n = 3) and severe cases (n = 6). ***Padj < 0.001 (Wilcoxon test, Padj was performed using Bonferroni correction).

**Figure 4.**
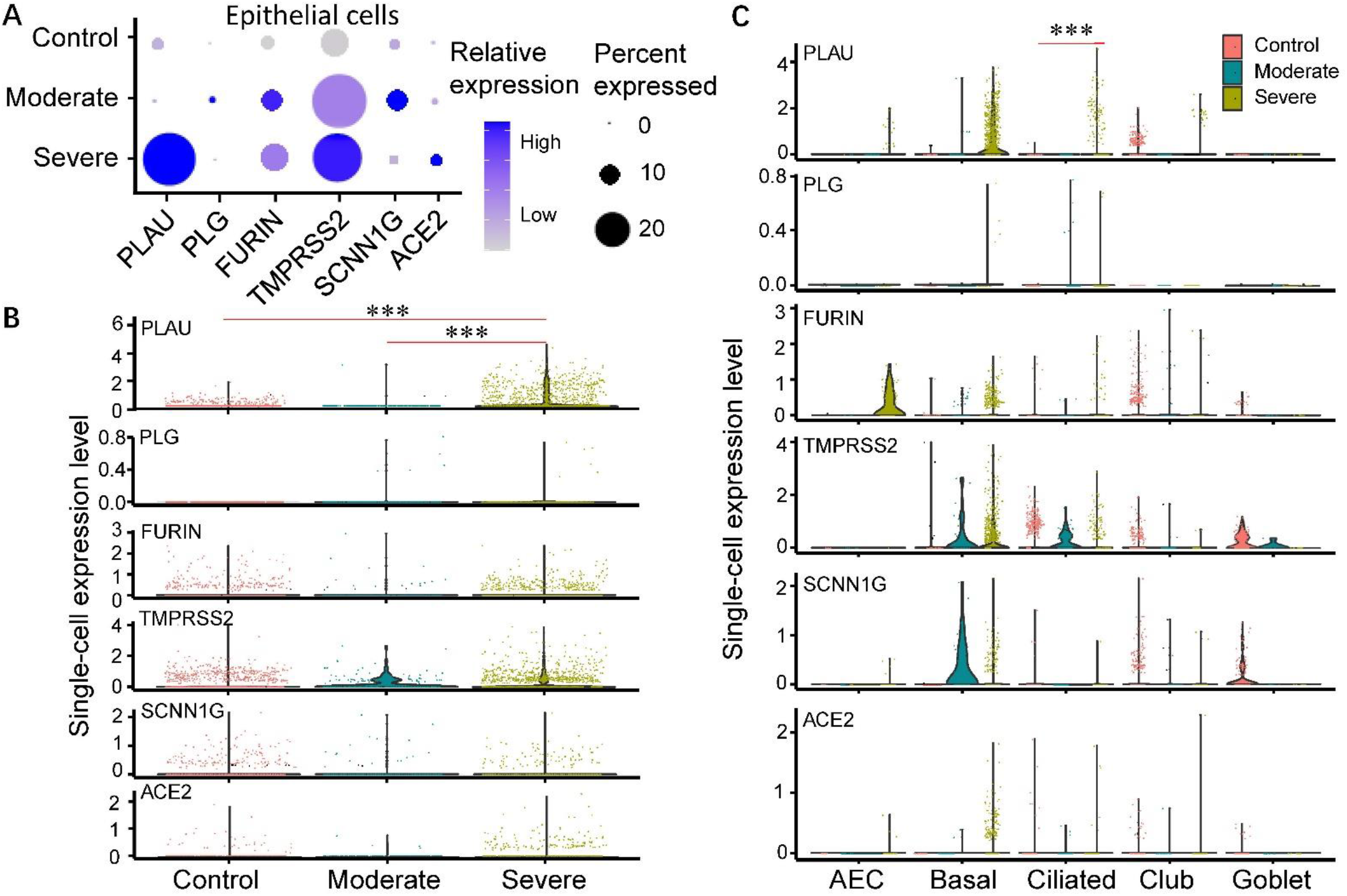
Transcription levels of proteases, ACE2, and SCNN1G in single epithelial cells of COVID-19 patients. **(A)** Bubble plot of SARS-CoV-2 receptor (ACE2) and proteases in BALFs epithelial cells of COVID-19 patients. The size of the dots indicated the proportion of cells in the respective cell type having a greater-than-zero expression of these genes, while the color indicated the mean expression of these genes. **(B)** The gene expression levels of selected proteases and ACE2 in epithelial cells from health controls (n = 4), moderate (n = 3), and severe cases (n = 6). **(C)** The gene expression levels of selected proteases and ACE2 in different epithelial cell types from health controls, moderate and severe cases. ***Padj < 0.001 (Wilcoxon test, Padj was performed using Bonferroni correction). AEC: alveolar epithelial cells.

The increments in PLAU (alveolar epithelial cells, basal, and ciliated cells), PLG (basal cells), FURIN (alveolar epithelial cells, basal, ciliated cells), TMPRSS2 (basal and ciliated cells), SCNN1G (alveolar epithelial cells and basal cells), and ACE2 (alveolar epithelial cells, basal, and club) were predominately observed in different cells. Especially, a significant increase in PLAU expression was seen in ciliated cells, while the expression of measured genes showed a decline in COVID-19 goblets (**Fig. 4C**). In addition, similar changes of these genes in leukocytes were shown in **Supplementary Fig. 2**.

To corporate the results in COVID-19 patients, we analyzed bulk-seq data of 3 human respiratory epithelial cell lines infected with SARS-CoV-2: A549, Calu-3, and NHBE (Blanco-Melo et al. 2020). PLAU transcript was significantly upregulated in all three cell lines after SARS-CoV-2 infection (multiplicity of infection = 2) (**Fig. 5**, *P* < 0.001). However, TMPRSS2 was only upregulated in infected Calu-3 cells, evidenced by recent studies (*P* < 0.001) (Xu et al. 2020). Similar to those of SARS and MERS, the SARS-CoV-2 infection also increased the expression level of ACE2 in A549 cells (*P* < 0.05) (Smith et al. 2020). Although SARS-CoV-2 did not change the mRNA level of SCNN1G significantly in these cell lines as that for influenza virus, researchers are warned to pay more attention to the post-translational modification of γENaC (Hou et al. 2019b).

**Figure 5.**
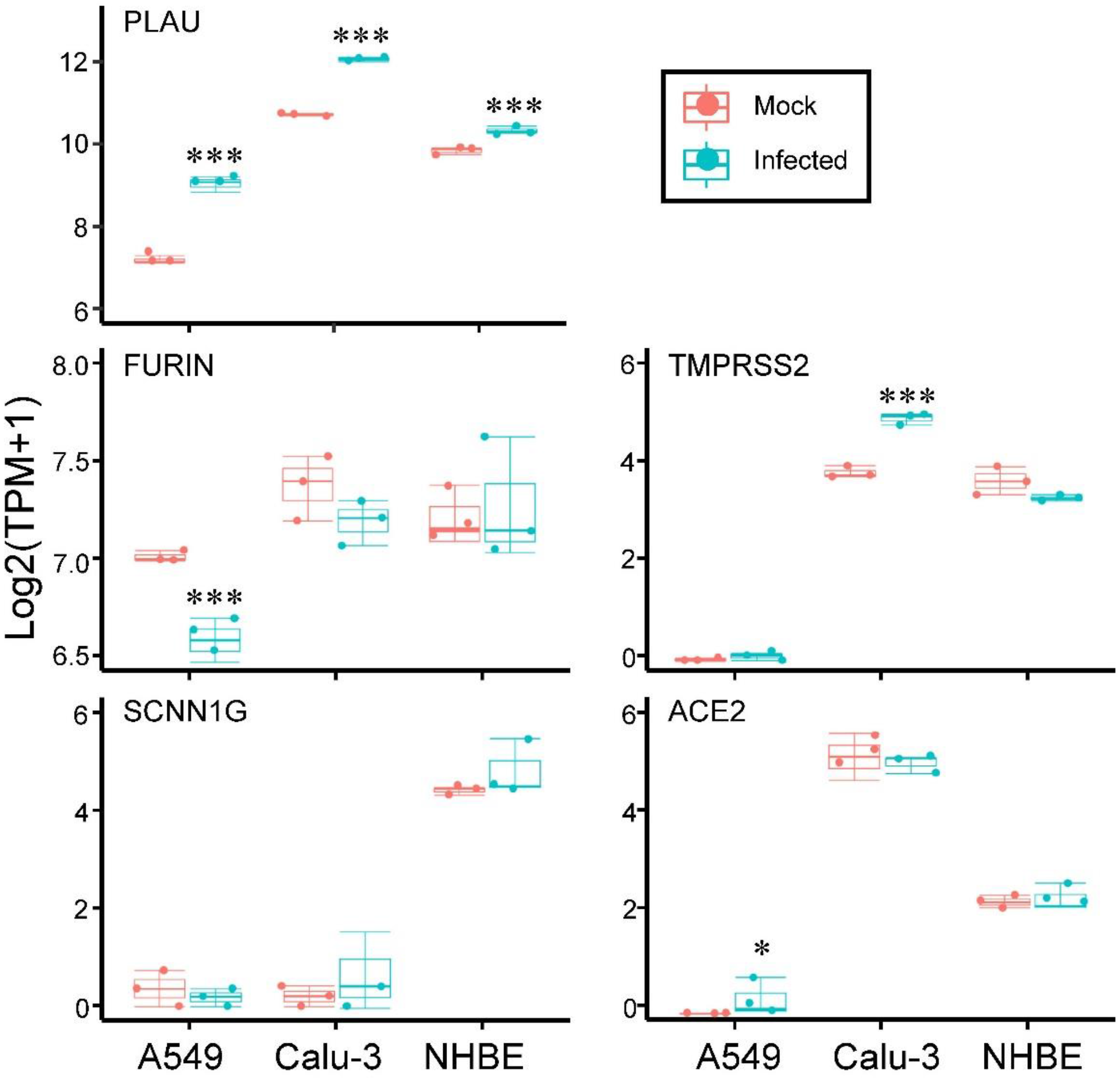
Changes of proteases, ACE2, and SCNN1G in respiratory cell lines after SARS-CoV-2 infection. Normal human bronchial epithelial (NHBE) and alveolar epithelial (A549, Calu-3) cells were infected with SARS-CoV-2 for 24 h (Infected), and control cells received culture medium only (Mock). The boxplot showed the changes of proteases (PLAU, FURIN, and TMPRSS), SCNN1G, and ACE2 in A549, Calu-3, and NHBE after SARS-CoV-2 infection. Differential genes were calculated by DESeq2, ***Padj < 0.001, *Padj < 0.05 (Wald test, Padj was performed using Benjamini-Hochberg post-hoc test).

## Discussion

The novel coronavirus, SARS-CoV-2, was identified as the causative agent for a series of atypical respiratory diseases, and the disease termed COVID-19 was officially declared a pandemic by the World Health Organization on March 11, 2020 (Pollard, Morran, and Nestor-Kalinoski 2020). SARS-CoV-2 has a great impact on human health all over the world, the virulence and pathogenicity of which may be relevant to the inserted furin site. Whilst the SARS-CoV-2 S2’ cleavage site has a similar sequence motif to SARS-CoV and would thus be suitable for cleavage by trypsin-like proteases, insertions of additional arginine residues at the SARS-CoV-2 S1/S2 (RRAR|S) clearly generate a furin cleavage site (Zhou et al. 2020). Interestingly, this difference has been implicated in the viral transmissibility of SARS-CoV-2 (Anand et al. 2020). Our data supported the investigation that furin sites (RRAR|S) not only exist in human virus but also in the γ-subunit of ENaC, which expresses highly in alveolar epithelial cells and a substrate to be cleaved by plasmin.

Plasmin has also been reported to have the ability to cleavage the furin site, and enhance the virulence and pathogenicity of viruses in their envelope proteins (Sidarta-Oliveira et al. 2020). SARS-CoV-2 has evolved a unique S1/S2 cleavage site, absent in any previous coronavirus sequenced, resulting in the striking mimicry of an identical furin-cleavable peptide on αENaC, a protein critical for the homeostasis of airway surface liquid (Anand et al. 2020). All the above indicates that SARS-CoV-2 infection will hijack the ENaC proteolytic network, which is associated with the edematous respiratory condition (**Fig. 6**) (Chen et al. 2014, Zhao, Ali, and Nie 2020). Our data showed that the respiratory cells co-express SARS-CoV-2 receptor, γENaC (SCNN1G), and plasmin family mainly belonged to alveolar type Ⅰ/Ⅱ, basal, club, and ciliated cells, respectively. The PLG (Plasminogen) expression in different cell types is not shown for its expression is too low to be detected in many lung scRNA-seq datasets. Of note, the ciliated cell is the predominant contributor to upregulate the PLAU gene in severe COVID-19 patients. As expected, PLAU levels, as well as TMPRSS2, are upregulated in respiratory epithelial cell lines after SARS-CoV-2 infection, supporting the idea that SARS-CoV-2 can facilitate ACE2-mediated viral entry *via* TMPRSS2 spike glycoprotein priming (Roberts et al. 2020). Enhanced PLAU expression induced by SARS-CoV-2 infection will activate the plasminogen, which may reduce the difficulty of SARS-CoV-2 invasion by cleaving the S protein.

**Figure 6.**
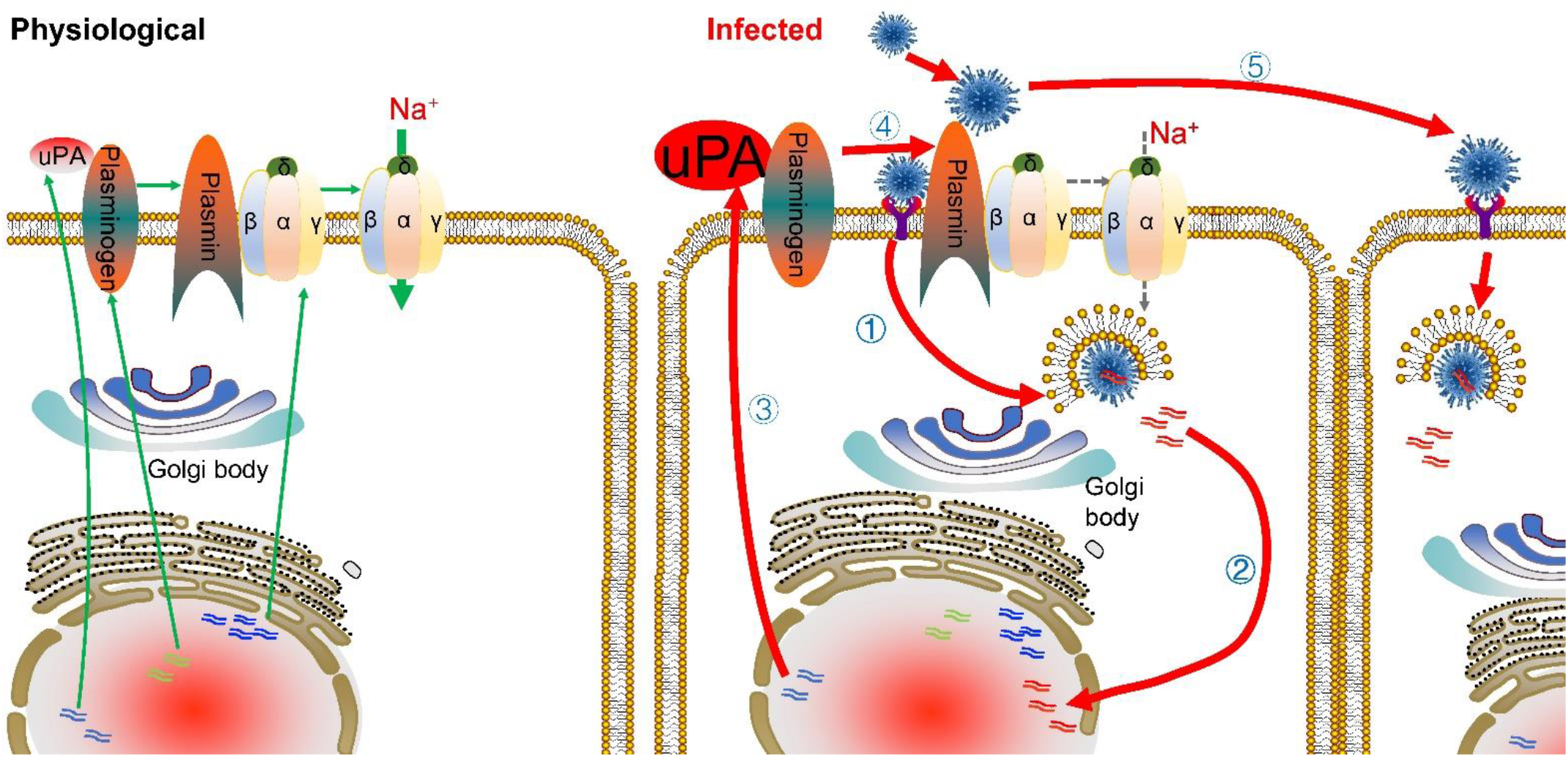
SARS-CoV-2 infection hijacks the ENaC proteolytic network. In physiological conditions, the urokinase activates the plasminogen to plasmin, which will cleave the γENaC, leading to its activation. After infected by SARS-CoV-2, the PLAU (urokinase) expression level is significantly upregulated, which may help other viruses’ invasion by activating the plasminogen to cleave the S protein. The green solid line represents the urokinase, plasminogen, ENaC mRNA transcripts and activation by plasmin under physiological conditions. The red solid line represents the activation process under infection conditions, while the grey dotted line denotes the repression effects.

The scRNA-seq data of bronchoalveolar lavage fluid cells from COVID-19 patients do not show the expression difference of SCNN1G (γENaC), which is considered to be regulated by plasmin through proteolytic hydrolysis. ENaC activity is not only determined by mRNA/protein expression but also cell proteases. Once the ENaC is biosynthesized and trafficked to the Golgi, it is likely to be modified by intracellular protease (furin). After inserted into plasma membrane, ENaC will encounter the opportunity for full proteolytic activation of the channel by extracellular proteases (elastase, plasmin, chymotrypsin, and trypsin) (Thibodeau and Butterworth 2013). Intriguingly, the PLG gene also did not show a difference between COVID-19 patients and healthy control, indicating that hyperfibrinolysis in COVID-19 patients may be induced by enhanced urokinase (Ji et al. 2020). Additional analysis of clinical studies or animal models is urgently needed to future explore the relationship between the plasmin, ENaC, and SARS-CoV-2 receptors at the protein level.

The amplified incidence of thrombotic events had been previously reported on COVID-19, and tissue plasminogen activator (tPA) was tried to treat stroke in COVID-19 patients (Vinayagam and Sattu 2020). We did not analyze the changes of PLAT in BALF cells of COVID-19 patients due to the tPA (PLAT) is generally expressed in endothelial cells. Similarly, the beneficial effects of plasmin on alveolar fluid clearance and novel mechanisms underlying the cleavage of human ENaCs at multiple sites by plasmin have been provided in our recent studies (Zhao, Ali, and Nie 2020). New drugs that regulate the uPA/ uPA receptor (uPAR) system have been demonstrated to help treat the severe complications of pandemic COVID-19 (D'Alonzo, De Fenza, and Pavone 2020). Amiloride, a prototypic inhibitor of ENaC, can be an ideal candidate for COVID-19 patients, supporting that ENaC is a downstream target of plasmin and involved in the luminal fluid absorption in SARS-CoV-2 infection (Adil, Narayanan, and Somanath 2020). Considering the two diametrically different therapeutic regimes in practice to address the complicated coagulopathic changes in COVID-19, fibrinolytic (alteplase, tPA) (Bona et al. 2020, Ly et al. 2020, Wang, Hajizadeh, et al. 2020, Barrett et al. 2020, Christie et al. 2020, Papamichalis et al. 2020, Poor et al. 2020, Arachchillage et al. 2020) and antifibrinolytic therapies (nafamostat and tranexamic acid) (Asakura and Ogawa 2020, Doi et al. 2020, Thierry 2020), our data provide new and comprehensive information on fibrinolytic related therapy targeting plasmin(ogen) as a promising approach to combat COVID-19.

## Methods

### Alignment of furin sites in viral and γENaC proteins

The sequences of γENaC proteins (rat, mouse, and humans) and human viruses were acquired from the UniProt (https://www.uniprot.org/). The accession numbers were P0DTC2 (for SARS-CoV-2), P04578 (HIV), P03435 (H3N2), A0A3G2XEB3 (Ebola), A0A140AYZ5 (MERS), P03188 (Epstein-Barr), P04488 (Herpes), P17763 (Dengue), P26662 (hepatitis), Q9Q6P4 (West Nile), A0A024B7W1 (Zika), P03420 (respiratory syncytial virus), P35253 (Marburg), P36334 (HCoV-OC43), A0A140H1H1 (HCoV-HKU1), P51170 (human γENaC), Q9WU39 (mouse γENaC), and P37091 (rat γENaC). Alignment was performed using the JalView software (Version: 2.11.1.0). The 3D structure of SARS-CoV-2 S (PDB ID: 6X2A) and γENaC (PDB ID: 6BQN) was modified and downloaded from the Protein Data Bank (http://www.rcsb.org/).

### Co-expression profiles of γENaC, ACE2, and proteases

We performed a systematic expression profiling of ACE2 and γENaC across 11 published human single-cell RNA sequence (scRNA-seq) studies comprising ~0.4 million cells using the nferX Single-Cell platform (https://academia.nferx.com/) (Anand et al. 2020). The mean expression of PLAU, SCNN1G, and ACE2 in a given cell-population (mean CP10k) was Z-score normalized (to ensure the Standard deviation = 1 and mean ~ 0 for all the genes) to obtain relative expression profiles across all the samples. The expression of PLAU, SCNN1G, and ACE2 in the respiratory system were analyzed and graphed as heatmaps using R package *pheatmap*.

### Acquisition, filtering, and processing of scRNA-seq data

The dataset downloaded from the Gene Expression Omnibus was filtered for integration. Lung scRNA-seq dataset (8 healthy controls in GSE122960) were filtered by total number of reads (nreads > 1,000), number of detected genes (50 < ngenes < 7,500), and mitochondrial percentage (mito.pc < 0.2). BALF scRNA-seq dataset was composed of 3 healthy controls, 3 moderate and 6 severe COVID-19 patients in GSE145926, and 1 healthy control in GSM3660650. These datasets were filtered by total number of reads (nreads > 1,000), number of detected genes (20 < ngenes < 6,000), and mitochondrial percentage (mito.pc < 0.1). Finally, a filtered gene-barcode matrix of all samples was integrated with the Seurat v3 to remove batch effects across different donors as described previously (Stuart et al. 2019).

### Dimensionality reduction and clustering

The filtered gene-barcode matrix was first normalized using the ‘LogNormalize’ methods in Seurat v.3 with default parameters. The top 2,000 variable genes were then identified using the ‘vst’ method in Seurat FindVariableFeatures function. Principal Component Analysis (PCA) was performed using the top 2,000 variable genes. Then Uniform Manifold Approximation and Projection for Dimension Reduction (UMAP) or t-Distributed Stochastic Neighbor Embedding (tSNE) was performed on the top 50 principal components for visualizing the epithelial cells. Meanwhile, the graph-based clustering was performed on the PCA-reduced data for clustering analysis with Seurat v.3. The resolution was set to 0.6 and 0.15 for the lung and BALF datasets to obtain a finer result, respectively. The markers used for BALF cell annotation were shown by the bubble plot in **Supplementary Fig. 3**.

### Differentiation of gene expression levels

Differentiation of gene expression level in BALF cells among the healthy, moderate, and severe groups was achieved using the Wilcox in Seurat v.3 (FindMarkers function). Then, we divided BALF cells into epithelial cells and leukocytes and compared gene expression levels among their subgroups. Both epithelial and leukocytes were re-clustered to detect the differences in gene expression of all cell types between healthy controls and severe/moderate COVID-19 patients. Bulk-seq data (GSE147507) was analyzed for the differential genes in respiratory epithelial cell lines using the DESeq2 with Wald test and Benjamini-Hochberg post-hoc test (Blanco-Melo et al. 2020, Love, Huber, and Anders 2014). It was considered significant if *P* < 0.05.

## Acknowledgment

This study was supported by NSFC 81670010, NIH grants HL87017, HL095435, and HL134828, AHA Awards AHA14GRNT20130034 and AHA16GRNT30780002. We were grateful to Yunlai Zhou (Yangzhou University) and Congxi Zhang (Gene denovo) for their assistance on bioinformatics.

