## Supplementary information for "Fibrinolysis influences SARS-CoV-2 infection in ciliated cells"

**Supplementary table 1.** List of single-cell studies analyzed and incorporated into the nferX resource (**https://academia.nferx.com/**)

| Title | Technique | Tissue | Pubmed ID or url |
|---|---|---|---|
| Host-viral infection maps reveal signatures of severe COVID-19 patients | 10X | Lung - bronchoalveolar lavage fluid | https://www.sciencedirect.com/science/article/pii/S0092867420305687 |
| A single-cell atlas of the human healthy airways | 10X | Lung, nasal cavity | https://www.biorxiv.org/content/10.1101/2019.12.21.884759v1 |
| scRNA-seq assessment of the human lung, spleen, and esophagus tissue stability after cold preservation | 10X | Lung | PMID: 31892341 |
| SARS-CoV-2 receptor ACE2 and TMPRSS2 are predominantly expressed in a transient secretory cell type in subsegmental bronchial branches | 10X | Lung | PMID: 32246845 |
| Single-cell analysis of olfactory neurogenesis and differentiation in adult humans | 10X | Respiratory tract, nasal cavity | PMID: 32066986 |
| Construction of a human cell landscape at single-cell level | Microwell-Seq | Lung | https://www.nature.com/articles/s41586-020-2157-4 |
| Virus-inclusive single-cell RNA sequencing reveals the molecular signature of progression to severe dengue | 10X | Lung | PMID: 30530648 |
| A cellular census of human lungs identifies novel cell states in health and in asthma | Drop-Seq | Lung | PMID: 31209336 |
| Construction of a human cell landscape at single-cell level | Microwell-Seq | Trachea | https://www.nature.com/articles/s41586-020-2157-4 |
| A cellular census of human lungs identifies novel cell states in health and in asthma | 10X | Nasal cavity | PMID: 31209336 |
| SARS-CoV-2 receptor ACE2 is an interferon-stimulated gene in human airway epithelial cells and is detected in specific cell subsets across tissues | Seq-Well | Lung | PMID: 32413319 |

**Supplementary table 2.** Top ten ranked cell populations expressing PLAU, SCNN1G, and ACE2

| **PLAU** | **SCNN1G** | **ACE2** |
|---|---|---|
| Neutrophils | Ionocytes | Club cells |
| Basal cells | AT1 | Mucous cells |
| Dendritic cells | Secretory cells | Ciliated cells |
| Monocytes | Club cells | Suprabasal cells |
| Fibroblasts | NK cells | AT2 |
| Club cells | T cells | Neutrophils |
| Macrophages | Endothelial cells | Deuterosomal cells |
| Endothelial cells | Suprabasal cells | NR4A1+ epithelial cells |
| Smooth muscle cells | Basal cells | Fibroblasts |
| Mast cells | AT2 | Smooth muscle cells |

**Supplementary figure 1.**

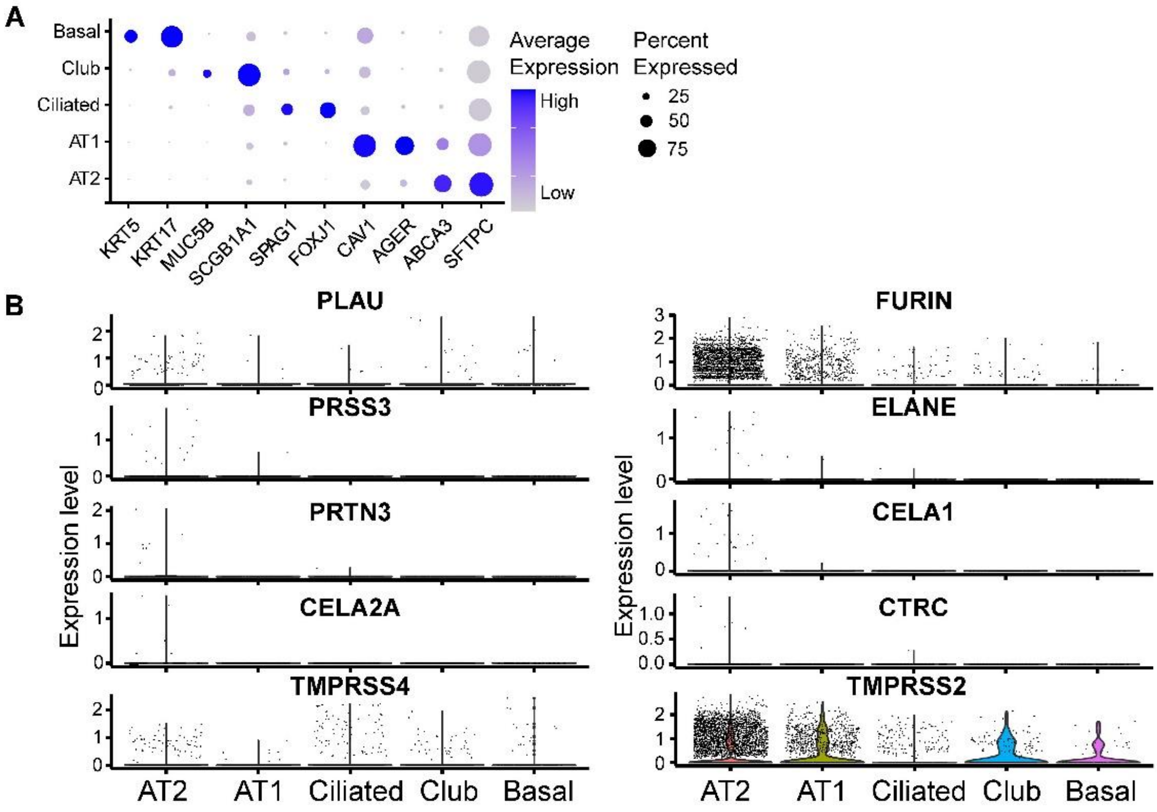


**Supplementary figure 1. Transcriptional levels of host proteases in respiratory epithelial cells. (A)** Bubble plot showed the epithelial cell markers used for cell annotation. **(B)** Violin plots showed the expression of urokinase (PLAU), furin (FURIN), trypsin (PRSS3), elastase (ELANE), proteinase 3 (PRT3), chymotrypsin-like elastase family member 1 and 2A (CELA1, CELA2A), chymotrypsin-C (CTRC), and transmembrane protease serine 2 and 4 (TMPRSS2, TMPRSS4) at the single-cell level.

**Supplementary figure 2.**

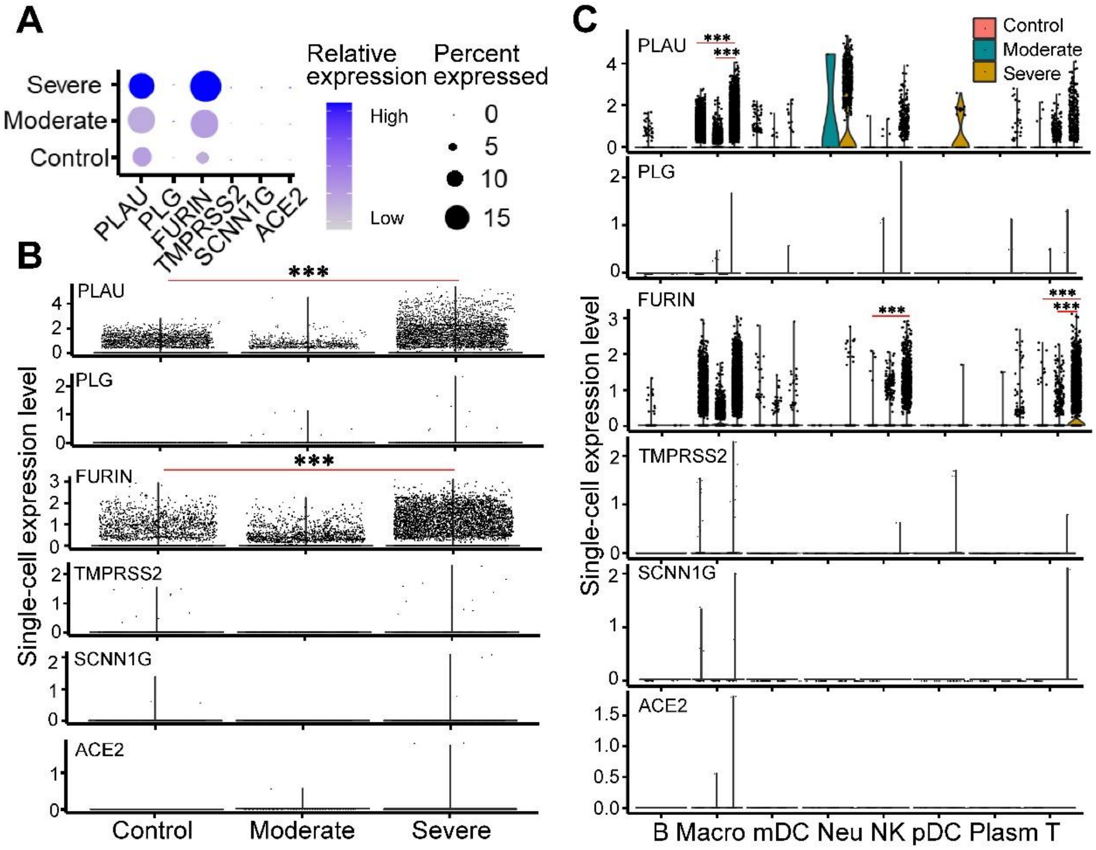


**Supplementary figure 2. RNA level of ACE2 and proteases in leukocytes of COVID-19 patients. (A)** Bubble plot of SARS-CoV-2 receptor (ACE2), and proteases in BALFs leukocytes of COVID-19 patients. The size of the dots indicateed the proportion of cells in the respective cell type having greater-than-zero expression of these genes, while the color indicateed the mean expression of these genes. **(B)** The gene expression levels of selected proteases and ACE2 in leukocytes from health controls (n = 4), moderate cases (n = 3) and severe/critical cases (n = 6). **(C)** The gene expression levels of selected proteases and ACE2 in different immune cell types from health controls (n = 4), moderate cases (n = 3) and severe/critical cases (n = 6). ***Padj < 0.001 (wilcoxon test, Padj was performed using bonferroni correction). Macro: macrophages, mDC: myeloid dendritic cells, pDC: plasmacytoid dendritic cells, Neu: neutrophils, NK: natural killer.

**Supplementary figure 3.**

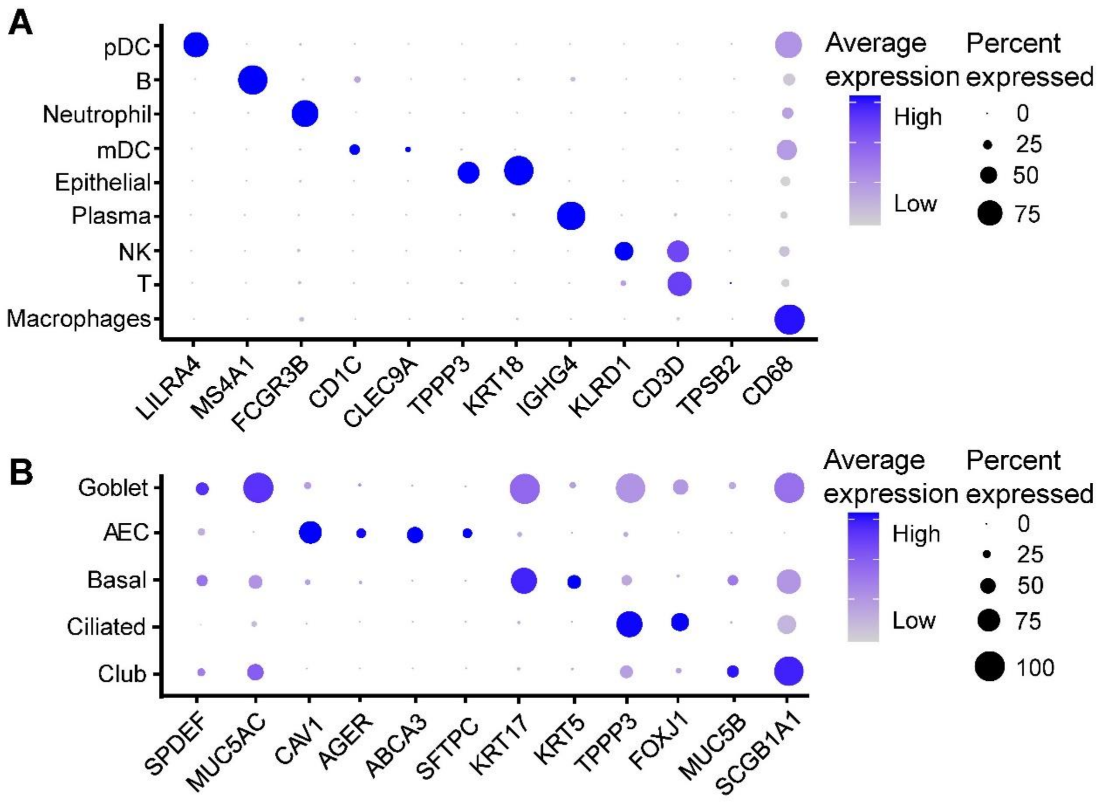


**Supplementary figure 3. Markers used for cell type annotation. (A)** Bronchoalveolar lavage fluid cell markers used for cell annotation. **(B)** Cell markers of epithelial cells in bronchoalveolar lavage fluid.
